# Promotion of tumor angiogenesis and growth induced by low-dose antineoplastic agents *via* bone-marrow-derived cells

**DOI:** 10.1101/2023.08.22.554227

**Authors:** Huining You, Peipei Zhao, Xue Zhao, Kai Cheng, Min Li, Jianrong Kou, Weiyi Feng

## Abstract

More research is needed to solidify the basis for reasonable metronomic chemotherapy regimens due to the inconsistent clinical outcomes from studies on metronomic chemotherapy with antineoplastic agents, along with signs of a nonlinear dose–response relationship at low doses. The present study therefore explored the dose–response relationships of representative antineoplastic agents in low dose ranges and their underlying mechanisms. Cyclophosphamide (CPA) and 5-fluorouracil (5-Fu) were employed to observe the effects of the frequent administration of low-dose antineoplastic agents on tumor growth, tumor angiogenesis, and bone-marrow-derived cell (BMDC) mobilization in mouse models. The effects of antineoplastic agents on tumor and endothelial cell functions with or without BMDCs were analyzed *in vitro*. Tumor growth and metastasis were significantly promoted after the administration of CPA or 5-Fu at certain low dose ranges, and were accompanied by enhanced tumor angiogenesis and proangiogenic factor expression in tumor tissues, increased proangiogenic BMDC release in the circulating blood, and augmented proangiogenic BMDC retention in tumor tissues. Low concentrations of CPA or 5-Fu were found to significantly promote tumor cell migration and invasion, and enhance BMDC adhesion to endothelial cells *in vitro*. These results suggest that there are risks in empirical metronomic chemotherapy using low-dose antineoplastic agents.

## Introduction

Despite studies on novel cancer treatments such as targeted therapies and immunotherapies are flooding the literature, clinical findings clearly indicate that chemotherapy still represents the current mainstay treatment strategy for cancer (1, 2). Standard clinical regimens for cancer chemotherapy typically employ the maximal tolerated dose (MTD) that has often been recommended as the regular or conventional model in the past, but often exerts only modest antitumor effects (3). This limited efficacy is in part due to the high toxicity of chemotherapeutic drugs to the host, which necessitates prolonged time intervals between treatment cycles to allow for normal tissue recovery from the cytotoxic assault (4). Moreover, cytotoxic agents in MTD chemotherapy decimate chemotherapy-sensitive cancer cell populations, leaving chemoresistant cells behind to recolonize the tumor bed, which ultimately leads to disease relapse between or after treatment cycles (5). This situation promoted the introduction of the new treatment strategy of low-dose metronomic (LDM) chemotherapy into clinical practice around 2000 (6). In contrast to MTD drug regimens, LDM chemotherapy is characterized by the administration of a cytotoxic agent at a lower and hence less-toxic dose at more-frequent regular time intervals, so as to optimize the antitumor efficacy and reduce the toxicity of antineoplastic drugs. LDM chemotherapy is expected to provide substantial benefits over MTD regimens by exerting both direct and indirect effects on tumor cells and the tumor microenvironment (TME) (7, 8). There is increasing evidence that LDM chemotherapy can inhibit tumor angiogenesis, stimulate the anticancer immune response, and induce tumor dormancy (9, 10).

Metronomic chemotherapy has been extensively reviewed in the literature and there is emerging evidence from preclinical and clinical studies that it has therapeutic benefits for both early- and advanced-stage malignant tumors. However, metronomic chemotherapy is still mainly used as a palliative care tool by most clinicians, rather than active, upfront therapy (11–13). Meanwhile, there have also been inconsistent and controversial findings about the efficacy of LDM chemotherapy (14–16). One of the important reasons for this dilemma may be that metronomic regimens of LDM chemotherapy have to be highly empirical due to the lack of a definitive method to optimize dosages, delivery schedules, and administration duration, despite the large number of clinical trials that have been performed (9, 17). LDM chemotherapy regimens currently applied in the clinic are actually mostly designed based on empirical evidence, and such treatments might not suppress tumor growth or even lead to chemotherapy failure if the dose and/or administration frequency are too low, and might induce serious side effects if these are too high. Dosage and treatment schedules of LDM chemotherapy in the clinic therefore usually have to be determined according to the dose–limiting toxicity experienced by patients or drug titration starting at a low dose for a period (e.g., 1 week), with the daily dose then increased each week to the point of maximum tolerance by the patient according to dose and frequency (18). The selection and judgment of the most-efficacious treatment is therefore subject to great uncertainty and is often delayed (19, 20). The development of metronomic regimens have therefore remained largely empirical, and there is no clear theoretical approach for the employed doses, administration frequency, and treatment duration.

It is also particularly noteworthy that there is evidence that cancer therapies can induce cellular and molecular responses in the tumor and host that allow them to escape therapy and promote progression (2, 21). For example, the pharmacological activities of some antineoplastic agents are opposite at low versus high doses *in vitro* and *in vivo*. This strange phenomenon has been observed for certain antitumor agents, with low-dose cyclophosphamide (CPA) found to facilitate tumor growth and tumor metastatic lung colonies in mouse models (22–24), and gemcitabine exhibiting dose–dependent biphasic effects on tumor growth in our previous experiments (25). These phenomena that some low-dose chemotherapeutic drugs promote tumor growth and metastasis were actually reported in a piecemeal fashion as early as the 1960s (26, 27). However, it is not clearly understood how the negative dose–response correlations are expressed in chemotherapy, which could provide clues for avoiding unintended effects in clinical metronomic chemotherapy.

There is emerging evidence that chemotherapy induces a rapid elevation of chemokine, cytokine, or growth factor levels followed by the rapid mobilization of protumorigenic cells that are recruited to the TME and lead to tumor growth (28). These host responses to therapy generate protumorigenic and prometastatic biological pathways, such as angiogenesis, tumor-cell epithelial-to-mesenchymal transmission (EMT) and cell proliferation, which partly explain treatment failure and acquired resistance (29, 30). Our previous study and others found chemotherapy-induced MMP-9 upregulation specifically in bone-marrow-derived cells (BMDCs), an effect which facilitates EMT in tumor cells and supports their growth by inducing angiogenesis, suppressing immune activities, or directly contributing to tumor resistance (31–33).

Myeloid-derived suppressor cells (MDSCs), a type of immunosuppressive BMDCs, are known as Gr-1^+^/CD11b^+^ myeloid cells in mice and have been associated with tumor growth and angiogenesis. Increased MDSC infiltration regulates tumor insensitivity to chemotherapy via the promotion of cytokine and proangiogenic factor secretion. These findings demonstrated that the delicate balance of BMDC activities in the TME is violated following tumor perturbation and treatment, further supporting the need for a better understanding of the complex relationship between chemotherapy and TME.

The nitrogen mustard alkylating agent CPA and the uracil fluorinated analog 5-fluorouracil (5-Fu) are commonly included in conventional and metronomic chemotherapy regimens, and palliative and adjuvant clinical treatments. Our previous research found that for CPA and some antineoplastic agents, the administration dosage was not positively correlated with tumor growth inhibition at low doses, and could even promote tumor growth at certain doses. However, few studies have focused on such a unique dose–response relationship or investigated the category ranges of antineoplastic agents for such phenomena, or the mechanisms underlying this novel pharmacological action in detail to determine the potential risks in chemotherapy under low-dose administration conditions. The DNA-alkylating agent CPA and antimetabolic agent 5-Fu, two cytotoxic antineoplastic agents that share nonidentical mechanisms, were therefore selected as the research objects in the present study to investigate dose–response relationships, response differences in tumor species, and their mechanisms in LDM chemotherapy for tumor growth and metastasis *in vitro* and *in vivo*.

## Method

### Animals and cell lines

Male C57BL/6 mice, KM mice and BALB/c mice (6-8 weeks) were purchased from the Experimental Animal Center of Xi’an jiao tong University. All animals were housed under constant temperature, humidity, and lighting (12h light per day) and allowed free access to food and water. All experiments were carried out in accordance with guidelines prescribed by the Ethics Committee at Xi’an jiao tong University. S180 sarcoma cells and Human umbilical vein endothelial cells (HUVECs) were kind gifts from Cancer Research Center of Xi’an Jiao Tong University. B16 melanoma and Lewis lung carcinoma (LLC) cells were obtained from National Key Urology Laboratory of First Affiliated Hospital of Xi’an Jiao Tong University. B16, LLC and HUVEC cells, were maintained in recommended medium supplemented with 10% FBS, 1% penicillin and streptomycin. All cell lines were cultured at 37°C in a humidified atmosphere of 5% CO_2_.

### Cell proliferation assay

MTT assay was used to assess cell proliferation, cell viability, and/or cytotoxicity. Briefly, 1 ×10^4^ HUVECs or B16 cells were plated in 96-well plates and cultured for 24h, and then antineoplastic agents-containing medium or CM (200 μL) were added for 48h. Cell proliferation assay was determined using a standard colorimetric MTT (3-4, 5-dimethylthiazol-2-yl-2, 5-diphenyltetrazolium bromide) assay. The absorbance of samples was then measured at a test wavelength of 490 nm.

### Cell migration and invasion assays

For migration assays, melanoma cells suspensions (200 μL, 2 × 10^4^ cells/well) were plated into the upper chamber of transwell plates (8 μm; Corning, USA), and the lower compartment was filled with 700 μL complete culture medium with corresponding concentrations of chemotherapy drugs. For invasion assays, 30 μL Matrigel Matrix (pre-diluted 1:5 with serum-free medium) (BD Biosciences, USA) was placed into the upper chamber of transwell plates, and B16 cells (200 μL, 4×10^4^ cells/well) were added 2 h later. Subsequently, 700 μL of the corresponding medium supplemented with 10% FBS was added into the lower chamber. After incubation for 24h at 37 °C, cells in the upper chamber were carefully removed. Migrated cells were fixed with 4% paraformaldehyde, stained with 0.1% crystal violet, and quantified using an inverted microscope.

### Cell adhesion assay

The 6 × 10^4^ BMDCs were resuspended with serum-free RPMI-1640 medium containing 2 μΜ Calcein-AM in a volume of 2 mL, and then placed in a cell culture incubator containing 5% CO_2_ at 37°C, stained for 30 minutes, and resuspended with RPMI-1640 medium containing 10% FBS. HUVECs were cultured in RPMI-1640 medium containing 10% FBS in a cell culture incubator containing 5% CO_2_ at 37°C, and then 2 × 10^4^ cells/well HUVECs were plated into 24-well plates. After 24 h, 0 μM BMDCs, 10 μM BMDCs conditioned medium were added to culture plate, and after 48 h, Calcein-AM stained BMDCs were added. Finally, we used microplate reader detecting fluorescence intensity and photographs were taken with a fluorescence microscope.

### Xenograft tumor model in vivo

Each mouse was injected subcutaneously with 1×10^6^ murine tumor cells (0.1 mL) in the right forelimb of the mice. The metastatic model was established by intravenously injecting tumor cells into the tail vein. CPA or 5-Fu was given intraperitoneally after tumor cells transplantation following chemotherapy schedules respectively. The control received vehicle only. Tumor size was measured using calipers, and tumor volume in each mouse was calculated based on the following formula: A×B^2^×0.5, where A and B are the larger and smaller diameter of the tumor, respectively. After a few days of treatment till tumor size reach to about 1 cm^3^, animals were euthanized by cervical displacement after anesthetization. Tumor tissues were resected, weighted, photographed and then fixed with 4% paraformaldehyde.

### Bone marrow transplantation

6 to 10-week-old recipient C57BL/6 mice were lethally irradiated (9 Gy) followed by bone marrow reconstitution by tail vein injection with 1×10^7^ BMDC cells isolated from green fluorescent protein-positive (GFP^+^) donor femurs. 8 weeks after bone marrow transplantation, the mice were used for tumor experiments.

### Histology and immunohistochemistry

Tissues were fixed with 4% paraformaldehyde and embedded in paraffin. Histological and immunohistochemical staining was performed using standard techniques as described previously. In brief, the tissue sections were stained immunohistochemically or with hematoxylin and eosin (H&E). The immunohistochemistry process was performed with Rabbit anti-CD31 (1:50 dilution, Abcam, USA), Rabbit anti-GFP antibody (1:500, AnaSpec, Fremont, USA) and Rabbit anti-laminin (1:1000, Sigma-Aldrich, Milwaukee, USA) as the primary antibody in accordance with the instructions provided with the Histostain-Plus kit (4Abio, China).

The intratumoral microvessel density (MVD) was determined on CD31 or laminin stained tumor tissues. Five fields were selected randomly from each tumor tissue section and quantitative analysis of the positively stained density was performed using Optimas image analyzer (Optimas Corporation USA).

### RT-PCR

Total RNA from mice tumors was extracted using TRIzol (Invitrogen) in accordance with the manufacturer’s instructions. The cDNA was synthesized via Prime Script RT Master Mix Perfect Real-Time kit (DRR036A; Takara, Tokyo, Japan). The primer sequences used in this study along with the expected product sizes are listed in Table 1. The cycling protocol for PCR involved incubating the samples at 94°C for 2 min followed by 35 cycles of denaturation at 94°C for 30 sec, annealing at 55°C for 30 sec, and extension at 72°C for 30 sec, with a final cycle of incubation at 72°C for 2 min. The amplification products were analyzed by electrophoresis (Beijing Junyi, Beijing, China) in agarose gels and detected under UV illumination (Bio-Rad Laboratories) after staining with nucleic acid dye (DuRed; FanBo Biochemicals, Beijing, China). Images were analyzed using a quantitative analysis system (Quantity One Analysis Software; Bio-Rad Laboratories).

### Flow cytometry

Cells isolated from peripheral blood were incubated with FITC anti-mouse CD11b antibody (BioLegend, 1:200), PE anti-mouse Gr-1 (BioLegend, 1:200), PE anti-mouse β3 (BioLegend, 1:150), PE anti-mouse CXCR4 (BioLegend, 1:80), or PE anti-mouse VEGFR2 (1:80, BioLegend) following a published protocol (34). The cells were subjected to flow cytometer on a FACScan (BD Bioscience, CA, USA) and data were analyzed with Cell Quest Software.

### Biological function and pathway enrichment analysis

KEGG and GO analyses of low-dose CPA and 5-Fu were carried out using Protein chip technology. GO analysis included the biological process, molecular function and cellular component.

### Statistical analysis

Data were represented as the mean ± SEM and determined by a two-tailed Student t test or one-way analysis of variance using the GraphPad Prism Version 8.0 software. P-values of 0.05 or less were regarded as statistically significant.

## Results

### Low-dose antineoplastic agents enhanced tumor growth *in vivo*

To investigate the impact of chemotherapeutic agents on tumor growth over a large dosage range, B16-melanoma-bearing C57BL/6 mice and S180-sarcoma-bearing KM mice were respectively treated using CPA at 2.5–40 and 10–80 mg/kg on alternating days for seven doses in total. As shown in Figure 1A–D, a dose–dependent biphasic response of B16 tumor growth to CPA was observed for the 2.5–40 mg/kg dose range, in which CPA treatment led to a marked increase of 107.3% in tumor weight (P<0.05) at 5 mg/kg and a significant reduction in tumor weight by 79.3% (P<0.05) at 40 mg/kg compared with the controls. A similar dose–dependent biphasic response on S180 tumor growth was obtained after CPA treatment. CPA significantly promoted tumor growth at 20 and 40 mg/kg by 72.6% and 117.5%, respectively (both P<0.01) (Figure S1A–D). CPA administered at 40 mg/kg via intraperitoneal injection every other day significantly enhanced Lewis lung carcinoma growth in a mice model 26 days after inoculation (Figure S1E&F).

**Figure 1.**
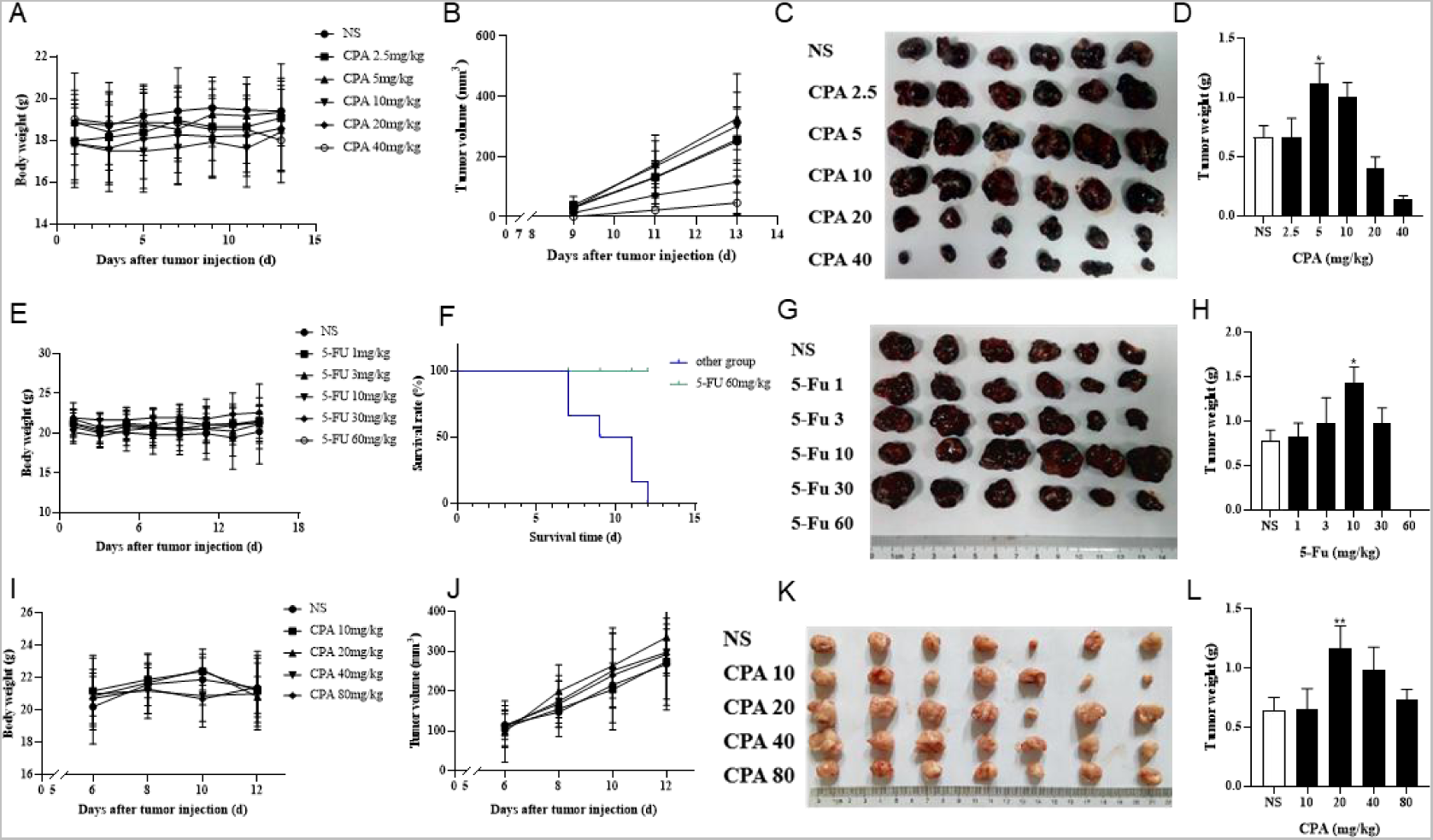
Low-dose CPA or 5-Fu promoted tumor growth *in vivo*. A-D, B16 tumor growth of mice treated with vehicle alone (normal saline) or the indicated dosages of CPA (n=6/group) A. Body weight; B. Tumor volume, monitored every two days; C. Macroscopic appearance of B16 tumors; D. Tumor weight; E-H, The survival curve and tumor growth of mice treated with vehicle alone or the indicated dosages of 5-Fu (n=6/group) E. Body weight; F. Survival curve; G. Macroscopic appearance of B16 tumors; H. Tumor weight; I-L, Growing S180 tumor growth of mice treated with vehicle alone or the indicated dosages of CPA (n=7/group) I. Body weight; J. Tumor volume; K. Macroscopic appearance of S180 tumors; L. Tumor weight. *vs* control, *, *P* < 0.05; **, *P* < 0.01.

The effects of 5-Fu on tumor growth were also tested in S180- and B16-bearing mice models. As shown in Figure 1E–H, 5-Fu injected at a low dose (10 mg/kg) on alternating days markedly increased B16 tumor weight by 84.8% (P<0.05) compared with the controls, while high-dose 5-Fu administration (60 mg/kg) resulted in the death of all six mice. Similarly, 5-Fu at 30 mg/kg critically promoted S180 tumor growth by 105.9% (P<0.01) in comparison with the controls (Figure S1G&H).

The effects of low-dose chemotherapeutic drug administration on growing tumor growth were also observed 6 days after tumor cell implantation (Figure 1I–L). S180 tumor growth was significantly increased by 81.7% (P<0.01) compared with the controls following the intraperitoneal injection of 20 mg/kg CPA. Similarly, 10 mg/kg CPA showed promoting effects of B16 tumor growth by 40.5% in comparison with the controls, but this change was not significant (P<0.06, Figure S1I–L).

CPA and 5-Fu could therefore promote tumor growth after being applied at certain low doses in a metronomic administration schedule.

### Low-dose antineoplastic agents facilitated tumor metastasis *in vivo*

The effects of low-dose CPA or 5-Fu on B16 tumor metastasis in lung tissues were investigated in mouse models. As shown in Figure 2A&B, 5 mg/kg CPA with an alternating-day administration schedule increased B16 tumor metastasis by 126.37%, and 40 mg/kg CPA induced a large decrease of 64.10% (*P*<0.05) in comparison with the controls. B16 tumor metastasis was significantly increased after treatment with 1 or 3 mg/kg 5-Fu every other day by 41.81% or 157.78% (*P*<0.05), respectively, and markedly decreased by 64.07% (*P*<0.05) with 30 mg/kg 5-Fu treatment (Figure 2C&D).

**Figure 2.**
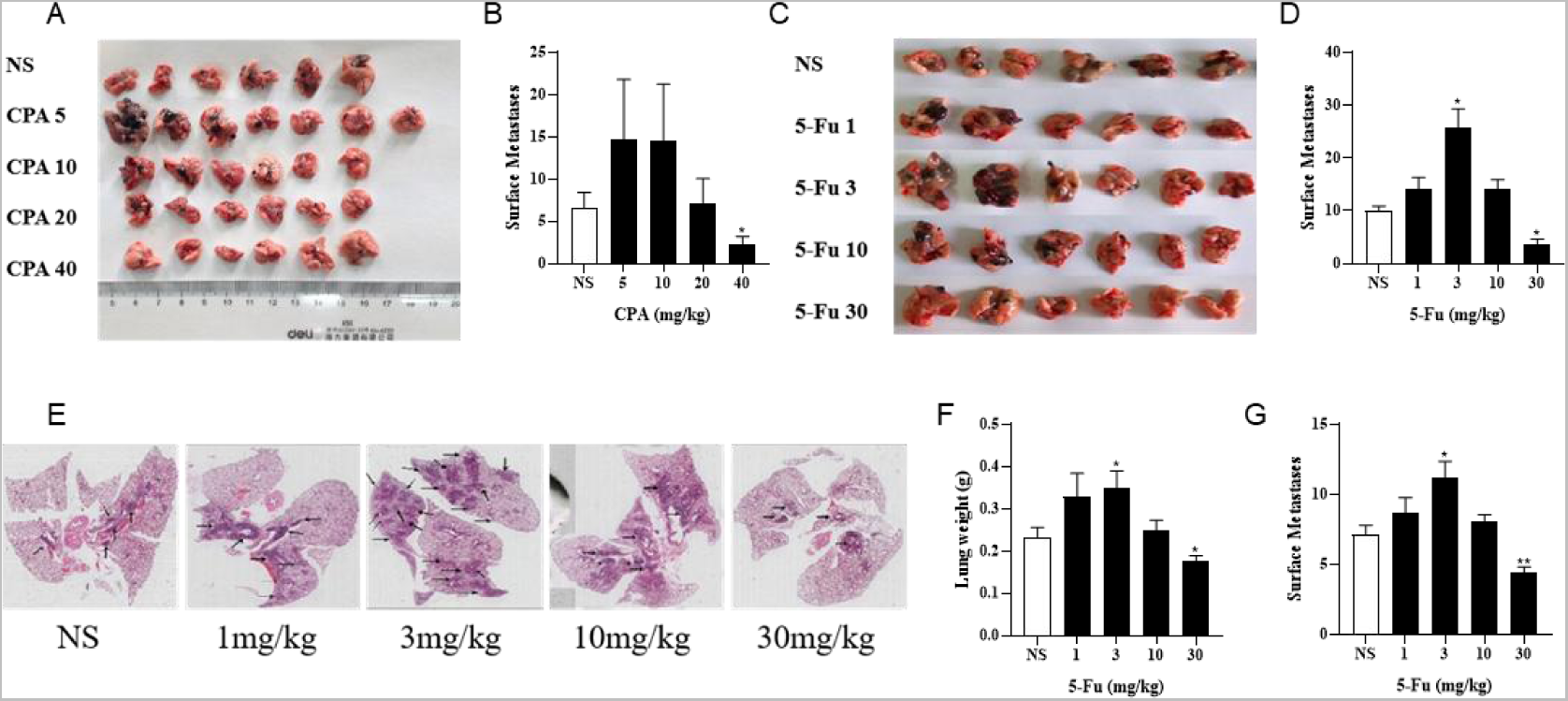
Low-dose CPA or 5-Fu facilitated tumor metastasis *in vivo*. A-B, B16 tumor metastasis of mice treated with vehicle alone (normal saline) or the indicated dosages of CPA (n=6-7/group) A. Effects of CPA on the metastasis of B16 tumor in mice; B. Surface metastases; C-D, B16 tumor metastasis of mice treated with vehicle alone or the indicated dosages of 5-Fu (n=6/group) C. Effects of 5-Fu on the metastasis of B16 tumor in mice; D. Surface metastases; E-G, LLC tumor metastasis of mice treated with vehicle alone or the indicated dosages of 5-Fu (n=6/group) E. Representative imges of the lung (Black arrow refers to tumor metastasis); F. Lung weight; G. Surface metastases. *vs* control, *, *P* < 0.05; **, *P* < 0.01.

As shown in Figure 2E–G, the metastatic tumor numbers and areas in lung tissue slides were obtained for 5-Fu-treated mice with Lewis lung carcinoma. The lung weights and metastatic tumor numbers in the group treated using 3 mg/kg 5-Fu were markedly increased by 65.2% and 56.1% (*P*<0.05), respectively, and those were significantly decreased in the group treated using 30 mg/kg 5-Fu by 21.7% and 38.7% (*P*<0.01) in comparison with the controls. Both CPA and 5-Fu therefore promoted tumor metastasis at certain low doses *in vivo*.

### Antineoplastic agents at low concentrations did not clearly accelerate tumor cell proliferation and viability *in vitro*

The effects of CPA and 5-Fu on B16 cell functions were evaluated to determine the role of tumor cells in chemotherapeutic-agent-induced tumor growth promotion. The results indicated that 48 h of CPA treatment led to the remarkable inhibition of B16 tumor cell proliferation in a concentration-dependent manner, for which the IC50 value of CPA was 158.1 μmol/L (Figure 3A&B). The viabilities of B16 tumor cells were also reduced after 48 h of incubation with low-dose CPA (Figure S2A&B). Similarly, 5-Fu treatment inhibited B16 cell proliferation in a concentration-dependent manner, with an

**Figure 3.**
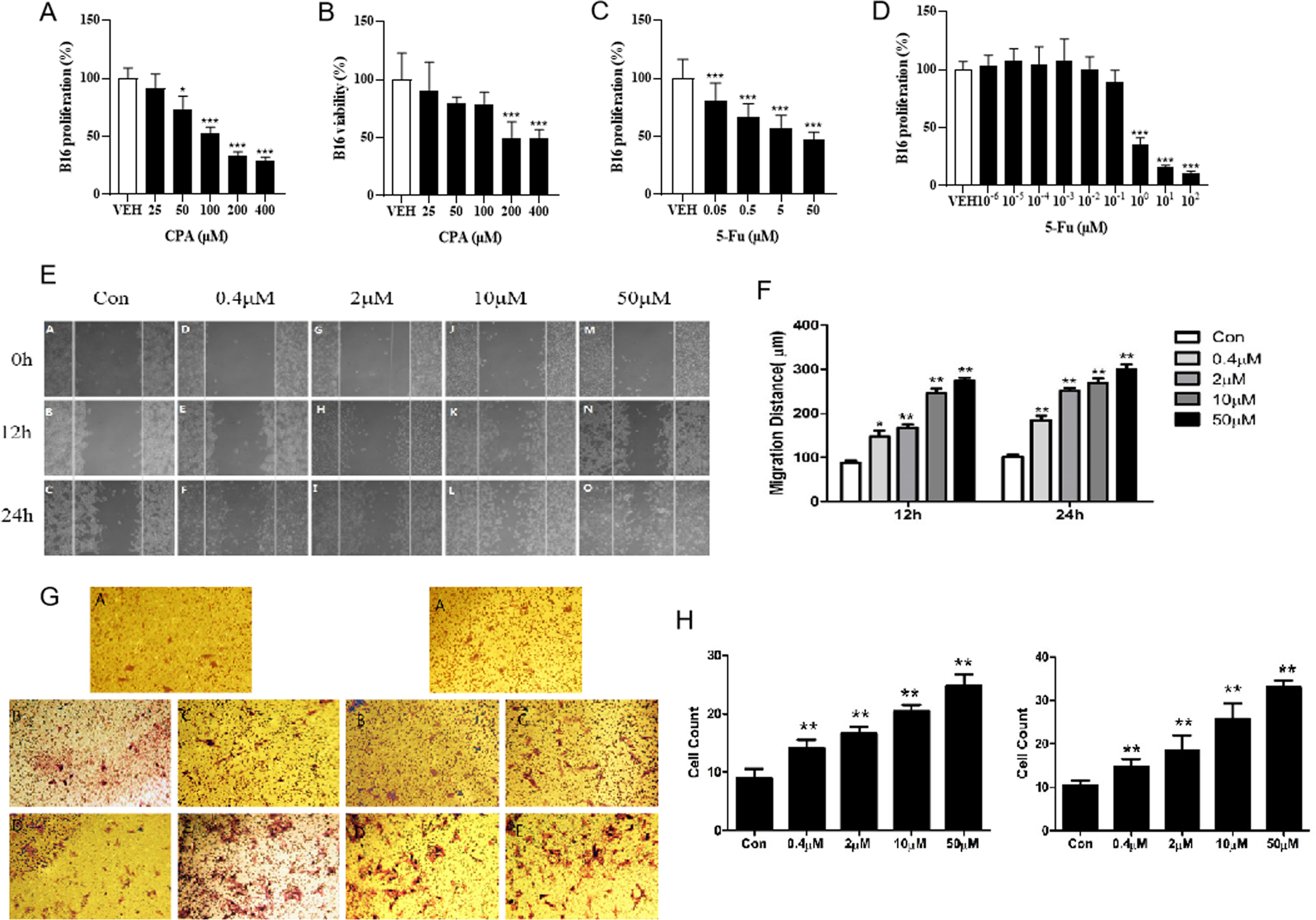
Low concentration of CPA or 5-Fu did not obviously accelerate tumor cells proliferation and viability *in vitro*, but significantly enhanced cell migration and invasion. A-B, The proliferation and viability of B16 treated with CPA (n=6/group) A. Cell proliferation; B. Cell viability; C. The proliferation of B16 treated with 5-Fu (n=6/group); D. The proliferation of B16 treated with low dose of 5-Fu (n=6/group); E. Representative images of scratch assay; F. Effects of 5-Fu on migration of B16 cells; G. Representative images of transwell experiment; H. Effects of 5-Fu on the migration and invasion of B16 cells. *vs* control, *, *P* < 0.05; **, *P* < 0.01, ***, *P* < 0.001.

IC_50_ value of 7.9 μmol/L (Figure 3C&D). Furthermore, the effects of 5-Fu on B16 tumor cell migration and invasion processes were evaluated using scratch-wound and Transwell assays, and tumor cell migration and invasion were promoted after 12 or 24 h of incubation at the concentration range from 0.4 to 50 μmol/L, respectively (Figure 3E–H). These results implied that tumor growth promotion induced by low-dose antineoplastic agents occurred via mechanisms including the promotion of tumor cell migration or invasion rather than by directly stimulating tumor cell proliferation.

### Low-dose antineoplastic agents promoted tumor angiogenesis *in vivo*

Since angiogenesis is necessary for solid tumor growth and metastasis, the effects of antineoplastic agents on tumor angiogenesis were analyzed using the laminin- and CD31-stained microvessel density (MVD). As shown in Figure 4A–E, the density of laminin^+^ vessels in 5 mg/kg CPA-treated B16 tumor tissues and 40 mg/kg CPA-treated S180 tumor tissues were significantly enhanced compared with the counterpart controls (both *P*<0.01), which was also the case in CD31^+^ vessels in 40 mg/kg CPA-treated LLC tumor models and 10 mg/kg 5-Fu-treated B16 tumor models (both *P*<0.05). Meanwhile, angiogenesis inhibition was observed in the high-dose administration group. These results imply that CPA and 5-Fu can promote tumor angiogenesis under continuous low-dose administration conditions and exert an inhibitory effect on angiogenesis at high doses.

**Figure 4.**
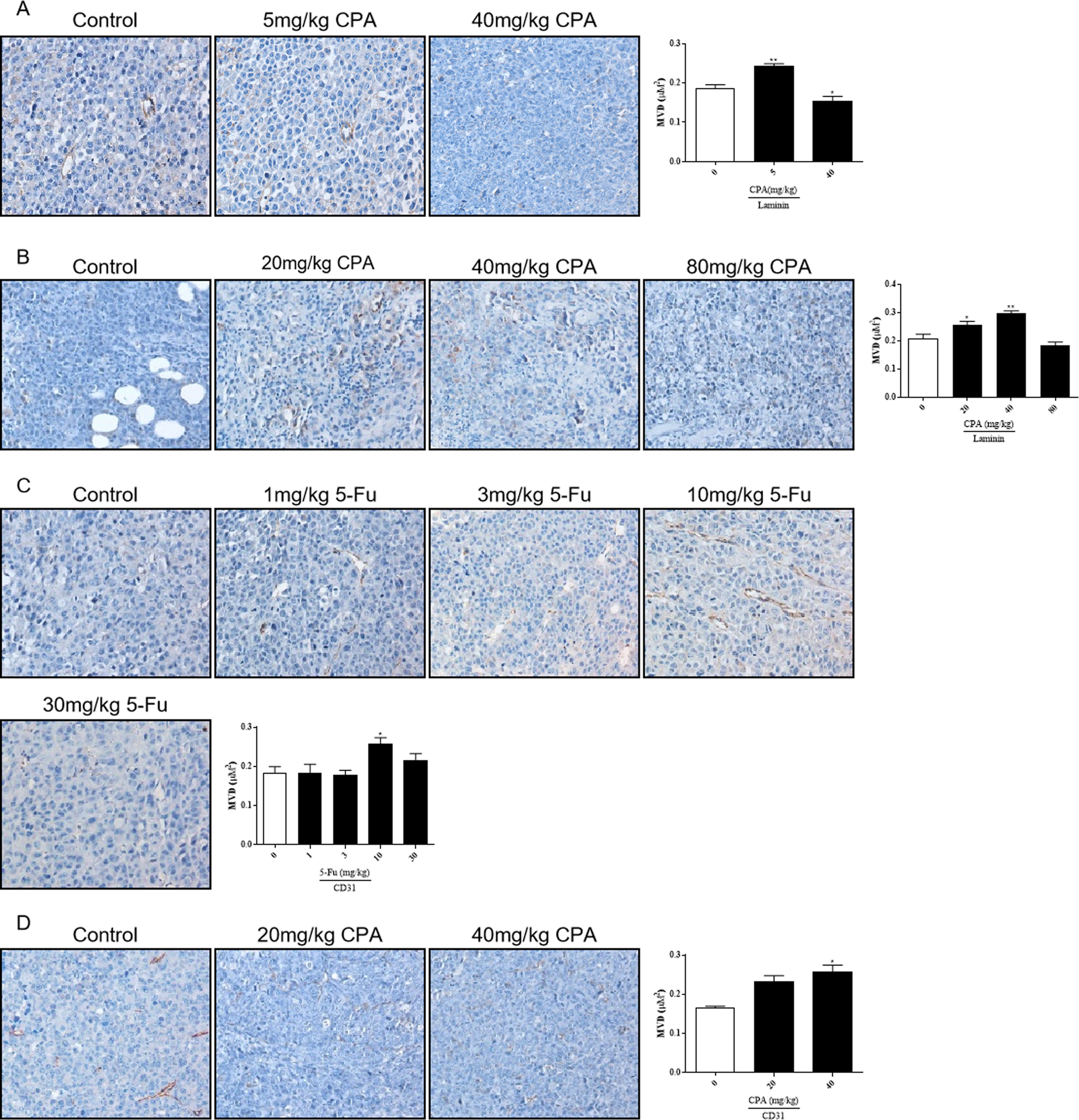
Effects of low-dose CPA or 5-Fu on tumor angiogenesis *in vivo*. A. Effects of CPA on micro-vessel of B16 tumor tissue (Laminin^+^); B. Effects of CPA on micro-vessel of S180 tumor tissue (Laminin^+^); C. Effects of 5-Fu on micro-vessel of B16 tumor tissue (CD31^+^); D. Effects of CPA on micro-vessel of LLC tumor tissue (CD31^+^). Scale bar, 20 μm, *vs* control, *, *P* < 0.05; **, *P* < 0.01.

A separate test was performed to compare the activities between CPA and 5-Fu and confirm these results (Figure S4A–D). B16 tumor growth was significantly promoted by 5 mg/kg CPA or 10 mg/kg 5-Fu treatments by 89.2% and 136.1%, respectively (*P*<0.05). Consistent with previous findings, 40 mg/kg CPA also inhibited tumor growth by 57.3% (*P*<0.05). Immunohistochemical staining results demonstrated significant increases in MVD in the groups that received 5 mg/kg CPA or 10 mg/kg 5-Fu, and increases in overall tumor necrosis in the high-dosage groups. These findings suggest that tumor angiogenesis was enhanced by low-dose CPA and 5-Fu (Figure S3E–H).

### Antineoplastic agents at low concentrations inhibited endothelial cell functions *in vitro*

Tumor angiogenesis depends on endothelial cell function. The effects of CPA or 5-Fu on endothelial cell functions were therefore investigated *in vitro*. Inhibitory effects of CPA on HUVEC proliferation (with an IC_50_ of 36.8 μmol/L) and viability were obtained, which were both significant when the concentration exceeded 10 μmol/L (Figure 5A&B). Similarly, HUVEC proliferation was observed to be significantly prohibited by 5-Fu at concentrations higher than 0.1 μmol/L, with an IC_50_ of 3.5 μmol/L, and viability was diminished at concentrations higher than 10 μmol/L *in vitro* (Figure 5C&D). However, the phenomena of promoting HUVEC proliferation and viability were not observed for either CPA at concentrations less than 10 μmol/L or 5-Fu at concentrations lower than 0.1 or 10 μmol/L *in vitro*, respectively (Figure 5E–H). These results indicated that both CPA and 5-Fu at low concentrations directly inhibited endothelial cell function *in vitro*.

**Figure 5.**
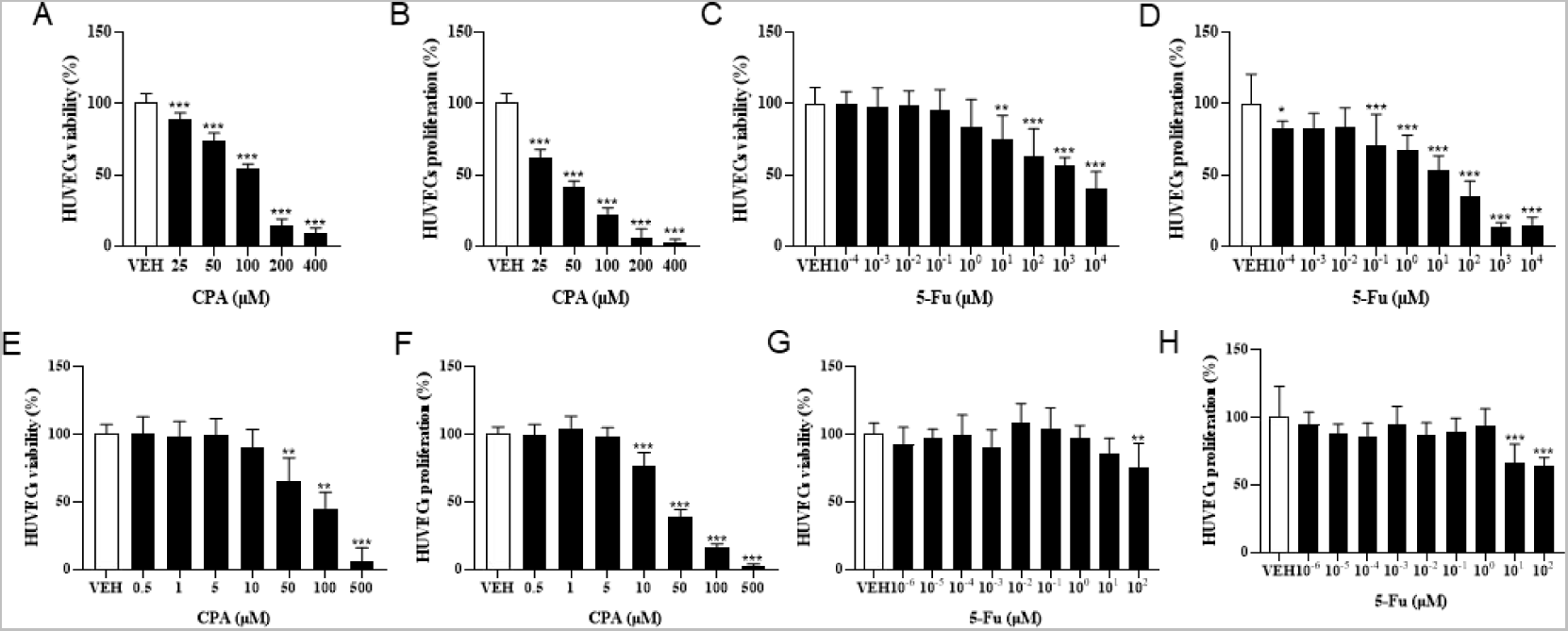
Low concentration of CPA or 5-Fu inhibited endothelial cell functions *in vitro*. A-B, The viability and proliferation of HUVECs treated with CPA (n=6/group); C-D, The viability and proliferation of HUVECs treated with 5-Fu (n=6/group); E-F, The viability and proliferation of HUVECs treated with low dose of CPA (n=6/group); G-H, The viability and proliferation of HUVECs treated with low dose of 5-Fu (n=6/group). *vs* control, *, *P* < 0.05; **, *P* < 0.01; ***, *P* < 0.001.

### Antineoplastic agents at low concentrations enhanced tumor and endothelial cell **functions through BMDCs**

To investigate the pathways of tumor growth and angiogenesis promotion induced by low-dose chemotherapeutic agent administration, the indirect effects of CPA or 5-Fu treatment on tumor or endothelial cell function were observed through BMDCs. As shown in Figure 6A&B, the proliferation rates were inhibited in a concentration-dependent manner in both B16 tumor cells and HUVECs after 48 h of CPA incubation *in vitro*. When treated using 100 and 200 μmol/L CPA, the proliferation was markedly decreased by 41.1% and 61.8% in B16 tumor cells, respectively, and by 29.9% and 31.8% in HUVECs compared with the counterpart controls (all *P*<0.001). In contrast, though no obvious effect of BMDCs conditioned medium was observed on B16 tumor cell or HUVECs proliferation, BMDC conditioned medium treated using 25 μmol/L CPA significantly enhanced HUVEC proliferation by 7.8% compared with a CPA-free BMDC conditioned medium (*P*<0.05).

**Figure 6.**
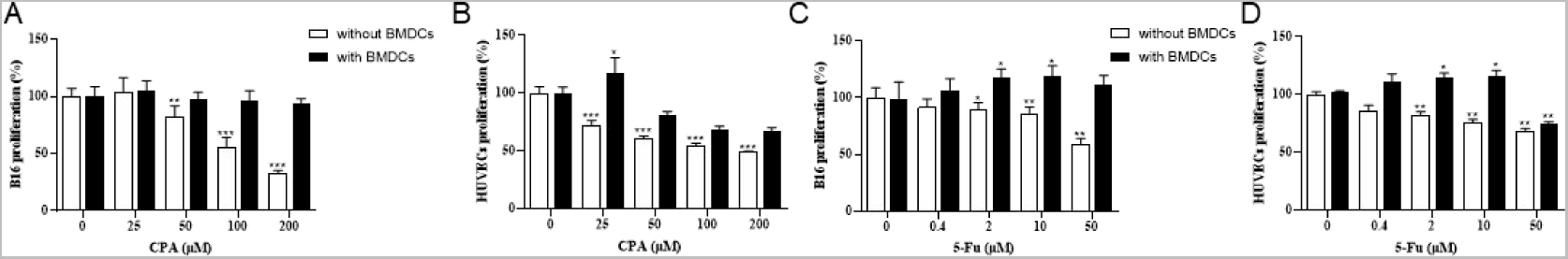
Low concentrations of CPA or 5-Fu promoted proliferation of tumor cells as well as endothelial cells through BMDCs. A-B, Effects of CPA-treated BMDCs conditioned medium on proliferation of B16 cells and HUVECs. (n=6/group); C-D, Effects of 5-Fu-treated BMDCs conditioned medium on proliferation of B16 cells and HUVECs. (n=6/group). *vs* control, *, *P* < 0.05; **, *P* < 0.01; ***, *P* < 0.001.

As shown in Figure 6C&D, 5-Fu exerted concentration-dependent inhibitory effects on B16 tumor cell and HUVEC proliferation rates. Compared with the counterpart controls, B16 tumor cell proliferation was significantly suppressed through 5-Fu administration at 2, 10, and 50 μmol/L by 8.3%, 12.5%, and 37.5%, respectively (*P*<0.05, *P*<0.01 and *P*<0.01), and HUVEC proliferation was also markedly suppressed by 15.0%, 21.7%, and 26.7%, respectively (all *P*<0.01). B16 tumor cell proliferation was significantly increased by 15.0% and 16.7% for 2 and 10 μmol/L 5-Fu, respectively (both *P*<0.05), and HUVEC proliferation was enhanced by 13.2% and 14.7% for 2 and 10 μmol/L 5-Fu, respectively (both *P*<0.05), and reduced by 22.1% for 50 μmol/L 5-Fu in comparison with the counterpart 5-Fu-free controls with BMDC conditioned medium (*P*<0.01).

These divergent results between with and without BMDC conditioned medium *in vitro* therefore imply that BMDCs might indirectly mediate the function enhancement of tumor and endothelial cells induced by antineoplastic agents at low concentrations.

### Antineoplastic agents promoting bone-marrow-derived proangiogenic cell release in the circulating blood and enhancing endothelial cell adhesion to BMDCs

Proangiogenic subtypes of BMDCs such as CD61^+^, Gr-1^+^CD11b^+^, VEGFR2^+^, and CXCR4^+^ in the circulating blood were analyzed in tumor-bearing mice using flow cytometry after CPA or 5-Fu treatment. As shown in Figure 7A–D, the counts of CD61^+^ BMDCs in the circulating blood of S180-bearing mice were significantly increased by 90.7% and 118.8% after treatment with 20 and 40 mg/kg CPA, respectively (both *P*<0.01), and by 71.5%, 58.9%, and 166.1% after treatment with 3, 10, and 30 mg/kg 5-Fu compared with the counterpart controls (*P*<0.05, *P*<0.05 and *P*<0.01, respectively). Meanwhile, Gr-1^+^CD11b^+^ BMDC release in the circulating blood was significantly enhanced by 54.4% (*P*<0.05) and 126.9% (*P*<0.001) at CPA dosages of 20 and 40 mg/kg, respectively, compared with the counterpart controls (Figure 7E&F). VEGFR2^+^ BMDC release was also significantly increased by 50.6% and 41.9% after CPA treatment at 20 and 40 mg/kg, respectively (*P*<0.01 and *P*<0.05, Figure 7G&H).

**Figure 7.**
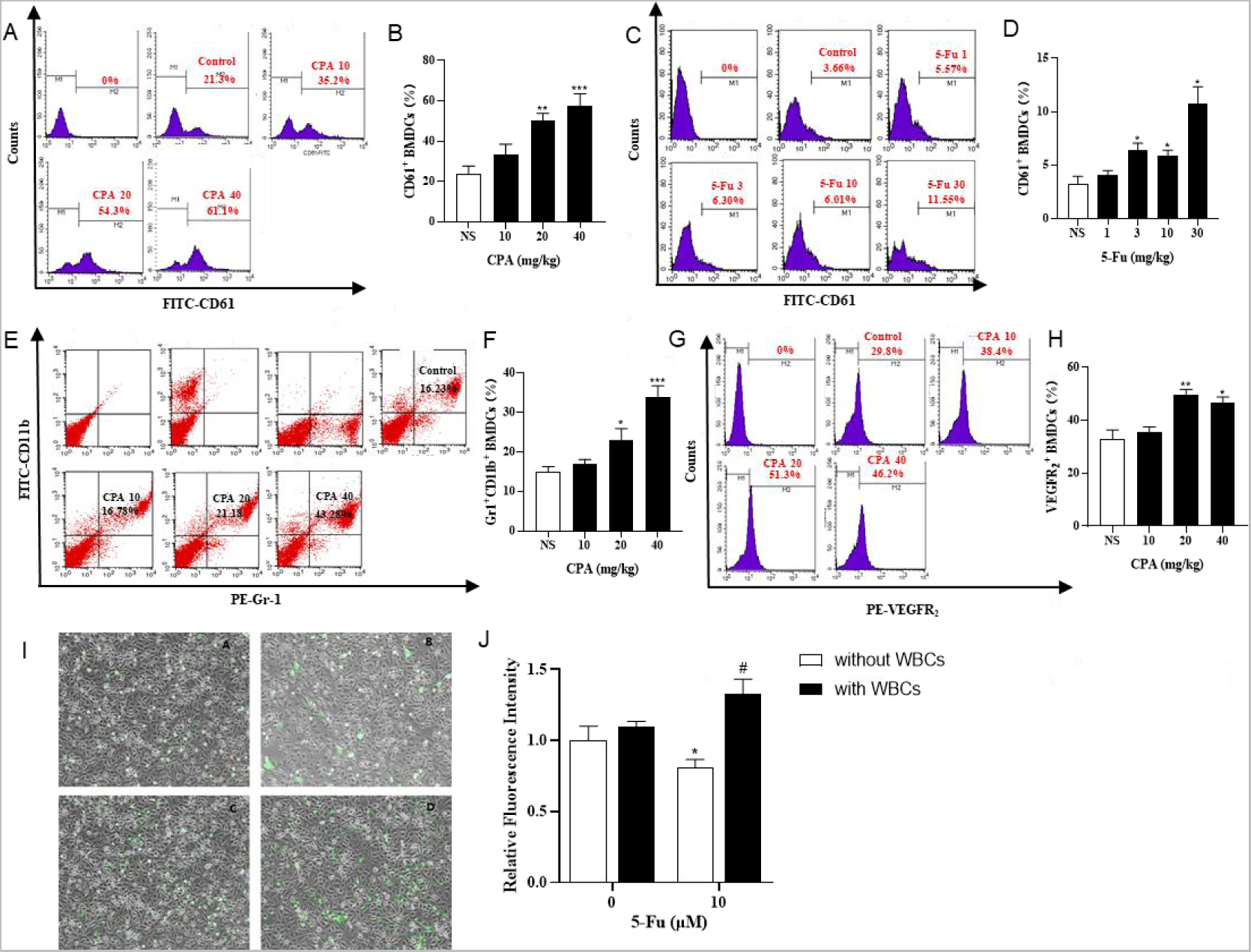
Chemotherapeutic drugs promoted BMDCs release in circulating blood and adhesion capacity of endothelial cells. A-D, CD61^+^ BMDCs counts in the circulating blood of S180 tumor-bearing mice treated with CPA and 5-Fu (n=5/group); E-F, Gr-1^+^CD11b^+^ BMDCs counts in the circulating blood of S180 tumor-bearing mice treated with CPA (n=5/group); G-H, VEGFR2^+^ BMDCs counts in the circulating blood of S180 tumor-bearing mice treated with CPA (n=5/group); I-J, Relative fluorescence intensity of HUVECs adhesion capacity treated with 10 μM 5-Fu or WBCs-conditioned medium (a: Control; b:10 μM 5-Fu; c: 0 μM 5-Fu/WBCs; d: 10 μM 5-Fu/WBCs) (n=5/group). *vs* control, *, *P* < 0.05; **, *P* < 0.01, ***, *P* < 0.001; vs the group of 0 μM 5-Fu treatment with WBCs, #, *P* < 0.05.

Endothelial cell adhesion is the main process of tumor metastasis, and adhesion capacity was investigated after 5-Fu treatment with or without BMDCs using fluorescence intensity indicators. The relative fluorescence intensity, which reflected the number of endothelial cells adhering to BMDCs, was 0.81±0.06 (mean±standard deviation) after 10 μM 5-Fu treatment, which was significantly lower than that of the controls (*P*<0.05). However, the adhesion effect of 10 μM 5-Fu with BMDCs on HUVECs was significantly enhanced, with a relative fluorescence intensity of 1.33±0.11 (*P*<0.05). These results implied that chemotherapeutic drugs promoted tumor growth and metastasis by inducing the promotion of BMDC release and adhesion to endothelial cells.

### Antineoplastic agents enhancing BMDC recruitment to tumor tissues and **promoting proangiogenic factors expression**

Tumor-bearing GFP^+^ bone-marrow-transplanted C57BL/6 mice were treated using 5 and 40 mg/kg CPA, and a dose–dependent biphasic effect on tumor growth was observed *in vivo*; GFP^+^ cell densities in B16 tumor tissues were then analyzed (Figure 8A–D). The results indicated that GFP^+^ cell densities increased by 198.9% (*P*<0.001) and 29.2% in the 5 and 40 mg/kg CPA groups, respectively. To explore the potential pathways underlying the bidirectional effects of antineoplastic agents on tumor growth, KEGG and GO analyses were conducted to analyze the enhancement or inhibition of tumor tissue growth induced by low- and high-dose chemotherapeutic drug administration using protein chip technology (Figure S4A&B). As a result, 13 pathways were enriched, of which the top 5 comprised the cytokine-cytokine receptor interaction, chemokine signaling pathway, intestinal immune network for IgA production, JAK-STAT signaling pathway, and autoimmune thyroid disease (Figure 8E).

**Figure 8.**
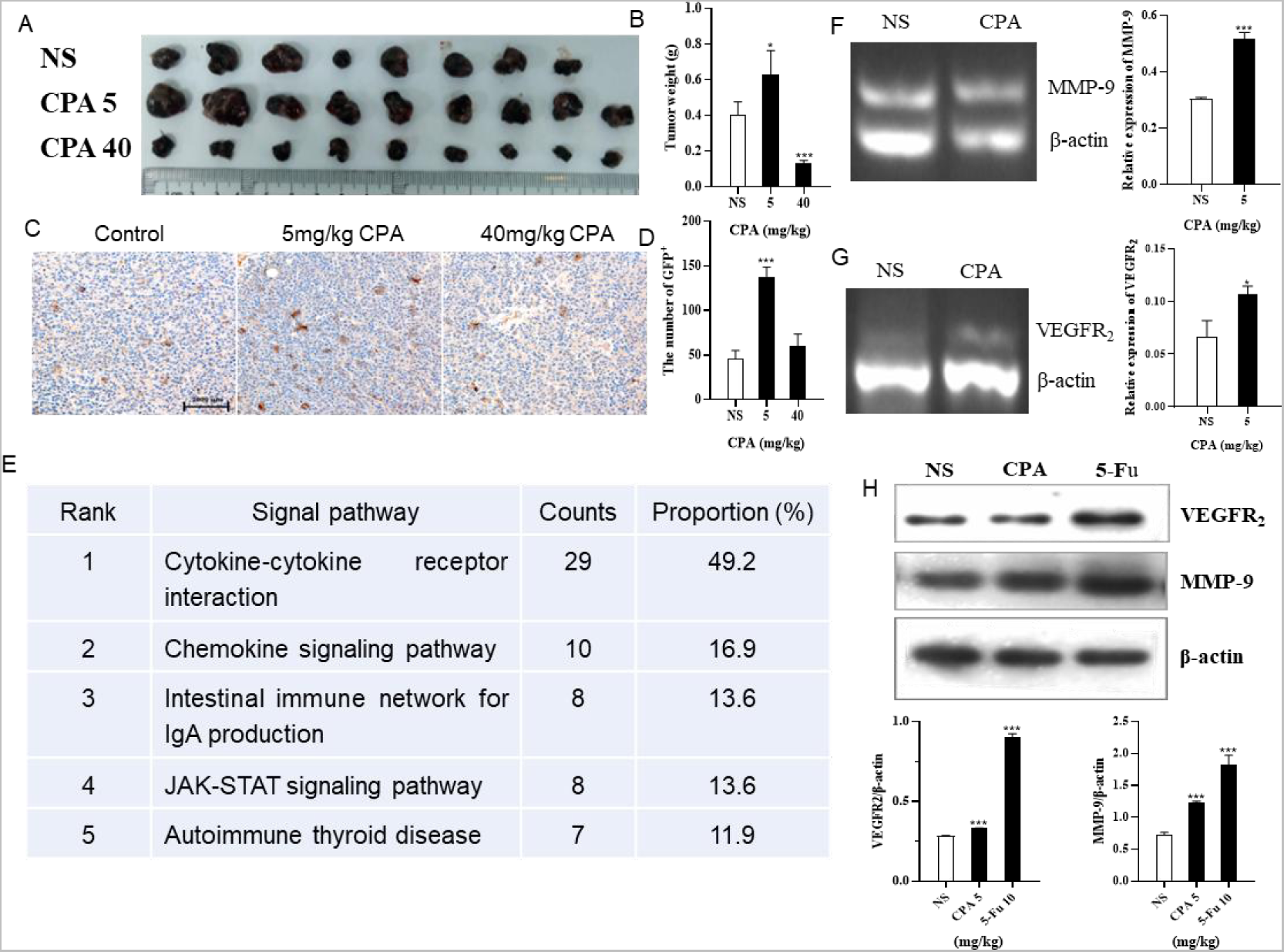
Low-dose CPA and 5-Fu promoted recruitment of BMDCs to tumor tissues and expression of pro-angiogenic factors. A-B, B16 tumor growth of GFP bone marrow-transplanted C57BL/6 mice treated with vehicle alone (normal saline) or the indicated dosages of CPA (n=8-9/group) A. Macroscopic appearance of B16 tumors; B. Tumor weight; C. IHC for GFP^+^ cells of B16 tumor tissues in bone marrow-transplanted mice (bar=2000 μm); D. The counts of GFP^+^ cells in tumor tissues; E. The top five KEGG pathways for the assemble caladium unigenes in control vs 5mg/kg CPA group; F-G, Effects of low-dose CPA on mRNA transcriptions of MMP-9, VEGFR2 in B16 tumor tissues (n=3/group) H. The expression levels of pro-angiogenic proteins, including MMP-9, VEGFR2. *vs* control, *, *P* < 0.05; ***, *P* < 0.001.

The transcription levels in tumor tissues of related proangiogenic cytokines or chemokines such as SDF-1, MMP-2, MMP-9, and VEGFR_2_ treated with CPA or 5-Fu were also determined using RT-PCR (Figure S4C&D). The results indicated that there were no significant differences in the transcription levels of SDF-1 and MMP-2 mRNA, but remarkable increases were observed in MMP-9 and VEGFR_2_ mRNA levels in the B16 tumor tissues of the 5 mg/kg CPA treatment group compared with the controls (both *P*<0.05,

). Similar results were obtained for 10 mg/kg 5-Fu-treated B16 tumor tissues (Figure S4E&F). As well as comparing the expressions of proangiogenic factors at the transcriptional level, their differences in protein expression were also compared. The results indicated that 5 mg/kg CPA and 10 mg/kg 5-Fu significantly improved MMP-9 expression levels by 69.4% and 152.8%, respectively, and enhanced VEGFR_2_ expression levels by 17.9% and 221.4% compared with the controls (Figure 8H).

These results suggest that low-dose CPA and 5-Fu promote GFP^+^ BMDC recruitment in tumor tissues and the expressions of proangiogenic factors and proteins.

## Discussion

The eradication of disease with relatively few side effects is the prime objective in clinical interventions. Due to frustration with conventional MTD outcomes, clinical oncologists are now increasingly resorting to the frequent, regular administration of LDM chemotherapy (35). Metronomic chemotherapy with continuous frequent administration of low-dose antineoplastic agents is novel in promoting antitumor efficacy with minimal toxicity while reducing the probability of acquired drug resistance developing. Metronomic chemotherapy has therefore recently been developed rapidly and has brought substantial benefits over MTD regimens (36). However, both the results of the present study and evidence from previous research have provided a new perspective regarding the effects of LDM chemotherapy on tumor growth, metastasis, and angiogenesis, in which some antineoplastic agents such as CPA and 5-Fu at certain doses in mice models were found to induce therapeutic outcomes that were contrary to treatment expectations. It was noteworthy in the present study that certain low doses of CPA and 5-Fu, which represent two antineoplastic agents with different pharmacological mechanisms, produced similar responses of promoting tumor growth in different tumor lines. To compare the effects of promoting doses of CPA with 5-Fu in S180 and B16 tumors with one dose administered every 2 days, CPA enhanced S180 tumor growth at low doses of 20 and 40 mg/kg and promoted B16 tumor growth at 5 mg/kg, while 5-Fu presented similar effects in facilitating tumor growth in S180 at 30 mg/kg and in B16 at 10 mg/kg. Low doses of other antineoplastic agents such as gemcitabine and cisplatin were also found to exhibit the same phenomenon of tumor growth enhancement in our previous studies (25). These results imply that it might be a common phenomenon for certain low doses of antineoplastic agents to exhibit a biphasic dose–response relationship and promote tumor growth.

The similar dose–response phenomenon of hormesis was proposed previously, where a toxicant would induce positive/stimulatory responses at low doses and negative/inhibitory responses at higher doses, therefore forming a biphasic dose–response relationship (37). These collective developments indicated that the hormetic dose–response model is the most common and fundamental in the biological and biomedical sciences, and is highly generalizable across biological models, endpoints measured, and chemical classes and physical agents (38). Studies have universally confirmed biphasic dose–response relationships similar to hormesis in 136 tumor cell lines for more than 30 tissue types and 120 different antitumor agents (39).

However, few previous studies have focused on the hormetic dose–response relationship between antineoplastic agents and tumor growth *in vivo*, although metronomic chemotherapy with low dosage is increasingly favored in clinics, especially in the context of metronomic chemotherapy regimens relying on empirical administration.

The data in the present study therefore confirmed the hormetic phenomenon of antineoplastic agents *in vivo* and hinted at the risks of empirical treatment regimens planning pattern, that is, antineoplastic agents may promote tumor growth once their concentration enters the low-dose stimulatory zone. To make matters worse, the unique nature of cancer treatment means that the treatment outcomes that promote tumor growth often go unnoticed. It is therefore strongly suggested that a precise method for dosing and timing selections for metronomic chemotherapy should be established based on sufficient experimental and theoretical evidence to avoid the risks of therapeutic outcomes that are inconsistent with or even opposing the treatment purposes. Unfortunately, no published studies have applied successful quantitative modeling to various possible clinical scenarios, nor mature theories to guide determination of the dosage and administration schedule. The dose range and amplitude of hormesis in antineoplastic agents are also still unclear, let alone its relationship with tumor species and drug classification.

In the present study, the dosages with promoting effects were usually 1/10∼1/2 of the inhibiting dose, which might partly overlap those in an empirical metronomic chemotherapy regimen, and stimulate tumor growth by 70∼100% relative to controls. In order to avoid promoting effects in an empirical metronomic chemotherapy protocol, it is therefore necessary to determine the threshold point and the stimulatory zone of antineoplastic agent doses, to elucidate the underlying mechanism of hormesis in antineoplastic agents, and to formulate a rational or optimum therapeutic regimen plan. The present results also indicated that 5 mg/kg CPA and 3 mg/kg 5-Fu promoted metastatic B16 melanoma or Lewis lung cancer cells in mice models, which further supports a hormetic dose–response relationship of antineoplastic agents in tumor metastasis. Several studies including our previous works have indicated that antineoplastic agents could promote cancer metastasis by directly influencing cancer migration and invasion, and indirectly establishing a favorable TME for cancer cell dissemination by modulating noncancerous cells (3, 25). There is therefore an urgent need to further investigate the mechanism and dose–response relationship of low-dose antineoplastic agents in tumor growth and metastasis.

While a dose–dependent bidirectional pharmacological action was observed in the present study, in which sustained low-dose chemotherapy significantly promoted tumor growth and metastasis while significantly inhibiting them at high doses, no direct stimulatory effects from antineoplastic agents on tumor cell proliferation were found at low concentrations *in vitro*, with even inhibitory effects being observed. These results suggest that the facilitated process of tumor growth at low doses was indirectly regulated rather than via direct stimulation of tumor cell proliferation by antineoplastic agents. At the same time, tumor cell migration and invasion were significantly increased by antineoplastic agent treatment at a proliferation-inhibiting low concentration, revealing that antineoplastic agents at low concentrations exert different effects on different functions of tumor cells.

Angiogenesis is crucial to tumor growth, progression, and metastasis (40). Based on the aforementioned previous research results, tumor angiogenesis was further analyzed *in vivo* and *in vitro* in the present study. The obtained data indicated that low doses of antineoplastic agents markedly promoted CD31^+^ and laminin^+^ tumor angiogenesis *in vivo* and inhibited the proliferation and viability of endothelial cell *in vitro*. These observations further suggest that low-dose antineoplastic agents act indirectly on the enhancement of tumor angiogenesis rather than by directly stimulating endothelial cell functions.

Our previous study found that BMDCs recruited by tumor tissues played a dominant role in promoting angiogenesis (33, 41). There is increasing evidence that antineoplastic agents actually stimulate the mobilization of proangiogenic BMDCs and their release from bone marrow, and enhance BMDC recruitment in tumor tissues (29, 42). The role of BMDCs in angiogenesis and tumor growth induced by low-dose antineoplastic agents were therefore investigated *in vivo* and *in vitro*. Our results indicate that in the presence of BMDCs, antineoplastic agents at the dosage that originally inhibited tumor and endothelial cell proliferation exerted stimulatory effects. This suggests that tumor angiogenesis and growth promoted by low-dose antineoplastic agents are mediated by BMDCs, and has revealed that BMDCs play a key role in this process.

Proangiogenic BMDC subtypes mobilized by antineoplastic agents in the circulating blood were then investigated, including β_3_^+^, VEGFR_2_^+^, and Gr-1^+^CD11b^+^ BMDCs, which were found at significantly increased levels due to the low-dose antineoplastic agents. The adhesive β_3_ integrin plays an important role in the process of BDMC adhesion, recruitment, and retention from the circulating blood to tumor tissues (33, 43). Studies have found that tumor angiogenesis was initiated by recruiting bone-marrow-derived VEGFR_2_^+^ endothelial progenitor cells, which differentiate into mature endothelial cells. VEGFR_2_ is considered the major mediator of proangiogenic signaling in almost all aspects of vascular-endothelial-cell biology, and mediates the proliferation, migration, and survival of endothelial cells (43). The findings that antineoplastic agents promote the mobilization of β_3_^+^, Gr-1^+^CD11b^+^, and VEGFR_2_^+^ BMDCs in the circulating blood and increase the recruitment and retention of GFP^+^ BMDCs in tumor tissues of bone-marrow-transplanted mice could therefore at least partly explain the phenomenon of tumor angiogenesis enhancement by LDM chemotherapy. Moreover, it is worth noting that proangiogenic Gr-1^+^CD11b^+^ BMDCs are also a common marker for immunosuppressive MDSCs. Researchers have paid considerable attention to the role of MDSCs in the TME that induce immunosuppression and immune escape by tumors (28). Antineoplastic agents have been found to increase the frequency and number of MDSCs in the circulating blood and tumor tissues of rodents and humans (44–46).Low-dose antineoplastic agents such as CPA and 5-Fu increase immunosuppressive proangiogenic Gr-1^+^CD11b^+^ BMDC levels, suggesting that they could promote tumor growth through the dual mechanism of enhancing angiogenesis and inhibiting immune function. The expressions of proangiogenic and immunomodulatory proteins in tumor tissues play an important role in tumor angiogenesis regulation, TME immune status, and the interplay between tumor-associated myeloid-cell-mediated immune suppression and angiogenesis (40, 47, 48).

The present study found significant increases in MMP-9 expression in mRNA and protein, the primary role of which is to degrade the extracellular matrix and enhance MDSC accumulation in the tumor tissues of groups with enhanced tumor growth. The expression of VEGFR_2_, a well-characterized receptor for vascular endothelial growth factor A, was also found to be markedly increased in the group that received LDM chemotherapy. However, there are distinct differences in protein expression caused by different antineoplastic agents, and the specific TME alteration caused by various chemotherapeutic drugs requires further study.

Together these findings indicate that while antineoplastic agents may partially kill dividing endothelial cells and probably also tumor cells, such drugs can directly mobilize and recruit proangiogenic BMDCs into local tumor tissues and support tumor growth upon therapy by enhancing angiogenesis. Although tumor cell proliferation and angiogenesis were directly inhibited by the cytotoxicity of antineoplastic agents, the strength of the effects of these agents in inhibiting endothelial or/and tumor cells and the drug-induced angiogenesis promotion and immunosuppression therefore determine whether tumor growth is enhanced or reduced. The present study has indicated that low-dose antineoplastic agents might not directly stimulate tumor or endothelial cell proliferation, but could indirectly intensify the mobilization and recruitment of BMDCs including MDSCs, upregulate the expressions of proangiogenic factors, and then facilitate tumor angiogenesis, which would result in the promotion of tumor growth and metastasis. Such a biphasic response from antineoplastic agents therefore needs more attention when selecting an appropriate dosage for achieving a desired pharmacological effect.

In conclusion, the results presented here indicate that LDM chemotherapy can promote tumor growth and metastasis, which is a common phenomenon in tumor treatment and mostly occurs due to the promotion of BMDC mobilization and recruitment, thereby fostering the expressions of various proangiogenic factors and tumor angiogenesis. Due to the potential risks associated with LDM chemotherapy, the optimal dosage and administration schedule of antineoplastic agents need to be determined through further research and theoretical support in order to achieve maximum therapeutic outcomes and avoid possible therapeutic risks from low-dose therapeutic regimens.

