## Supplemental figure and table for "Promotion of tumor angiogenesis and growth induced by low-dose antineoplastic agents *via* bone-marrow-derived cells"

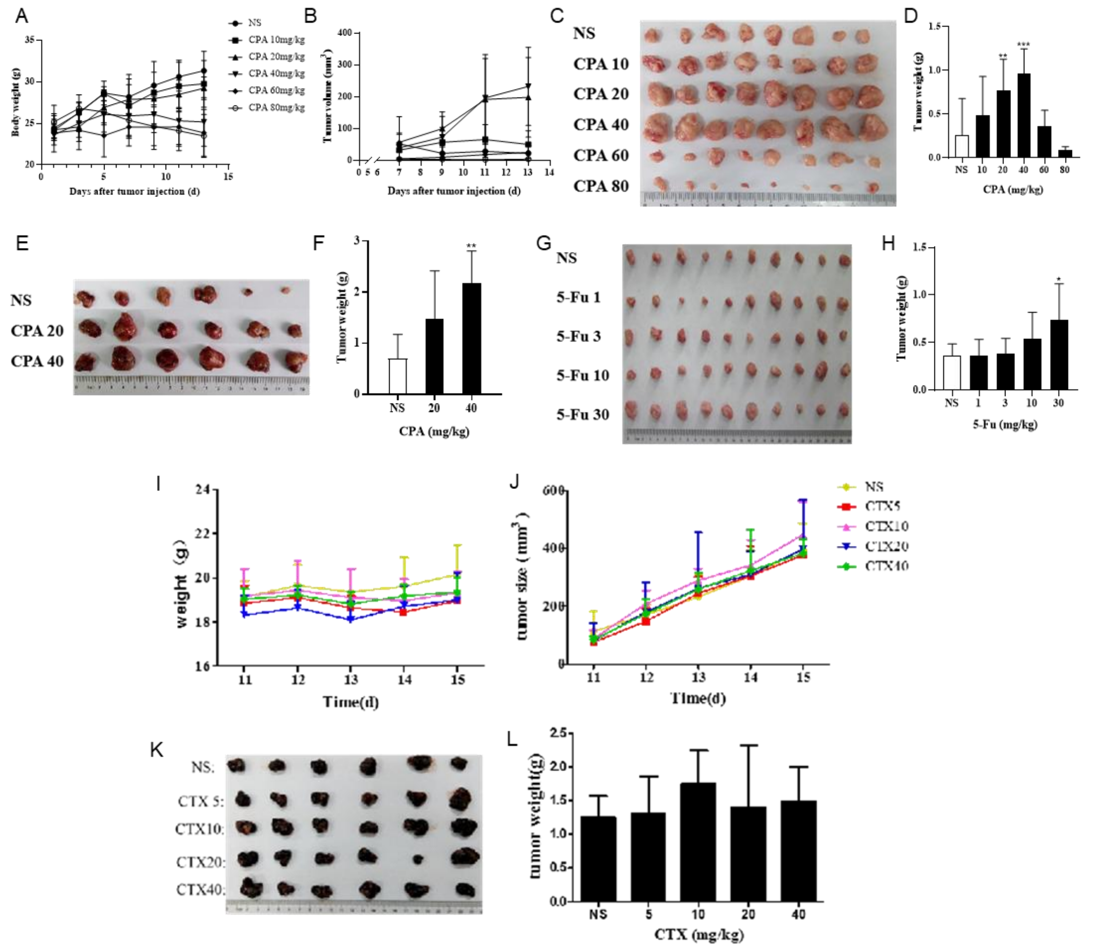


**Supplementary Figure 1 Low-dose antineoplastic agents promoted tumor growth in vivo.**

A-D, S180 tumor growth of mice treated with vehicle alone or the indicated dosages of CPA (n=8/group) A. Body weight; B. Tumor volume, monitored every two days; C. Macroscopic appearance of S180 tumors; D. Tumor weight; E-F, LLC tumor growth of mice treated with vehicle alone or the indicated dosages of CPA (n=6/group) E. Macroscopic appearance of LLC tumors; F. Tumor weight; G-H, S180 tumor growth of mice treated with vehicle alone or the indicated dosages of 5-Fu (n=10/group) G. Macroscopic appearance of S180 tumors; H. Tumor weight; I-L, Growing B16 tumor growth of mice treated with vehicle alone or the indicated dosages of CPA (n=6/group) I. Body weight; J. Tumor volume; K. Macroscopic appearance of B16 tumors; L. Tumor weight. vs control, *, $P < 0.05$; **, $P < 0.01$; ***, $P < 0.001$.

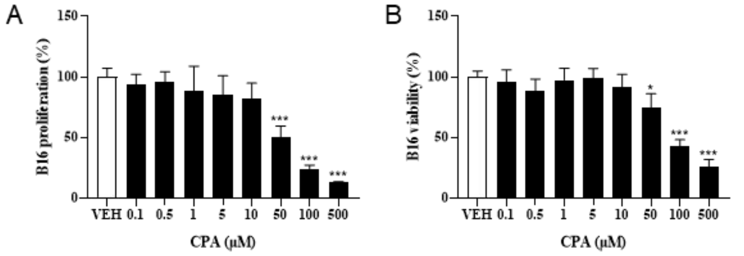


**Supplementary Figure 2 Low-dose CPA did not show promoting effects in tumor cells proliferation and viability in vitro.**

A. The proliferation of B16 treated with low dose of CPA (n=6/group); B. The viability of B16 treated with low dose of CPA (n=6/group). vs control, *, P < 0.05; ***, P < 0.001.

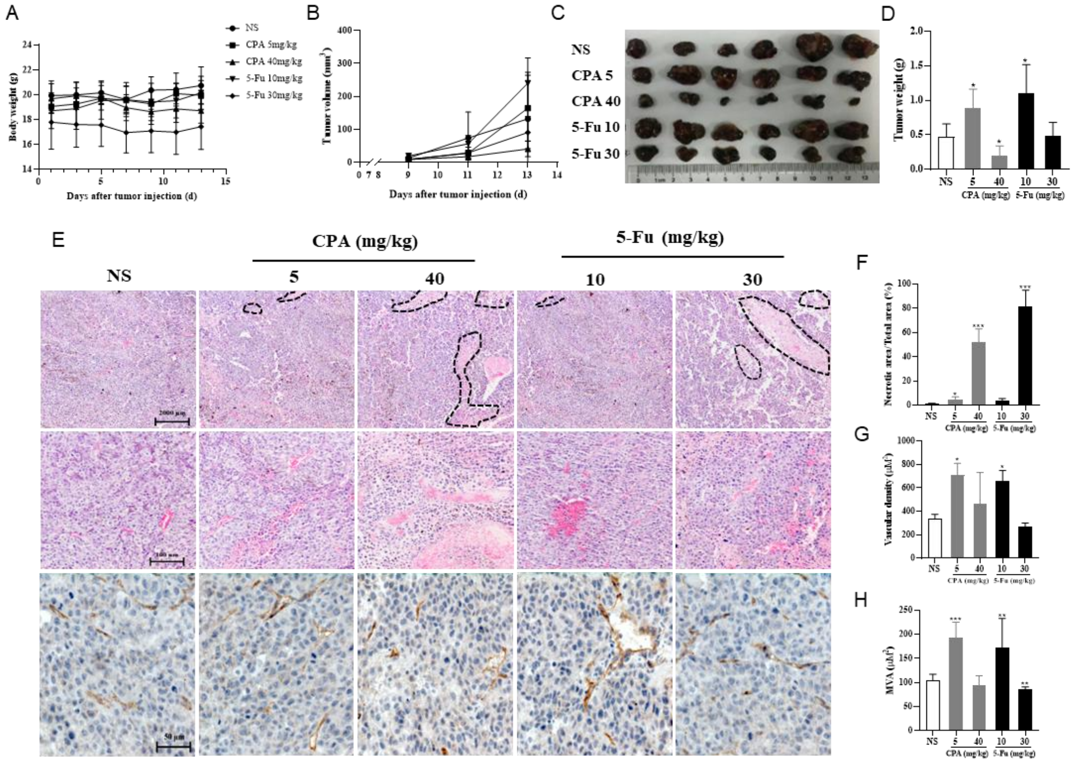


**Supplementary Figure 3 Comparison of anti-tumor efficacy and micro-vessel area between high-dose and low-dose CPA and 5-Fu treated B16 tumor models.**

A-D, B16 tumor growth of mice treated with vehicle alone (normal saline) or the indicated dosages of CPA and 5-Fu (n=6/group) A. Body weight; B. Tumor volume, monitored every two days; C. Macroscopic appearance of B16 tumors; D. Tumor weight; E. Tumor necrosis, tumor vessel and micro-vessel of B16 tumor tissues; F. Effects of CPA or 5-Fu on tumor necrosis of B16 tumor tissues; G. Effects of CPA or 5-Fu on vascular density of B16 tumor tissues; H. Effects of CPA or 5-Fu on micro-vessel area of B16 tumor tissues. vs control, *, P < 0.05; **, P < 0.01; ***, P < 0.001.

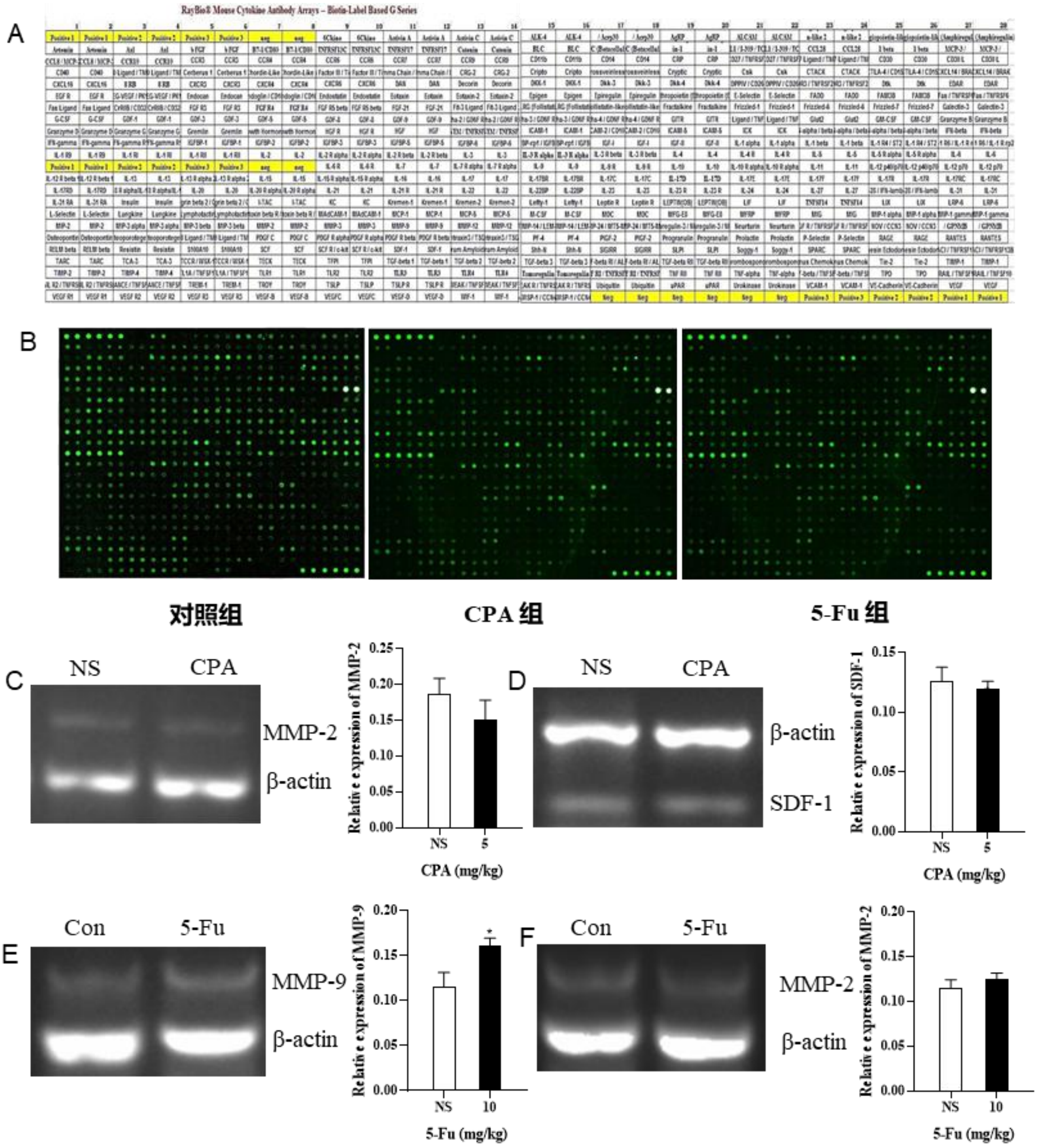


**Supplementary Figure 4 Low-dose CPA and 5-Fu promoted recruitment of BMDCs to tumor tissues and expression of pro-angiogenic factors.**

A. The protein chip assay; B. The protein expression patterns of B16 tumors from mice treated with vehicle alone, CPA and 5-Fu; C-D, Effects of low-dose CPA on mRNA transcriptions of MMP-2, SDF-1 in B16 tumor tissues (n=3/group); E-F, Effects of low-dose 5-Fu on mRNA transcriptions of MMP-9, MMP-2 in B16 tumor tissues (n=3/group). vs control, *, P < 0.05.

Table 1. Primer sequences for RT-PCR.

| Primers | Sequences (5'→3') | Product size (bp) |
|---|---|---|
| Mouse SDF-1 (F) | AGCCAACGTCAAGCATCTG | 106 |
| Mouse SDF-1 (R) | TAATTTCGGGTCAATGCACA | |
| Mouse β-actin (F) | GAGACCTTCAACACCCCAGC | 263 |
| Mouse β-actin (R) | ATGTCACGCACGATTTCCC | |
| Mouse $VEGFR_2$ (F) | CGCTCAGTGATGTAGAGGAAG | 343 |
| Mouse $VEGFR_2$ (R) | CAGAGCAACACACCGAAAGAC | |
| Mouse MMP-2 (F) | TAGACCTCAGCTTGCCCATT | 402 |
| Mouse MMP-2 (R) | CCTTGGTGGAACAGAAGGAA | |
| Mouse MMP-9 (F) | GGCACCTACTTGCTCACCTC | 106 |
| Mouse MMP-9 (R) | GTGTGTGTGTATGCCCAAGC | |
